# An optimized strategy for cloning-based locus-specific bisulfite sequencing PCR

**DOI:** 10.1101/239566

**Authors:** Mario Van Poucke, Xanthippe Boulougouris, Bart De Spiegeleer, Christian Burvenich, Luc Duchateau, Luc J. Peelman

## Abstract

In this methods paper, we describe a successful strategy to investigate locus-specific methylation by cloning-based bisulfite sequencing. We cover sample handling, DNA isolation, DNA quality control before bisulfite conversion, bisulfite conversion, DNA quality control after bisulfite conversion, *in silico* identification of CpG islands, methylation-independent bisulfite sequencing PCR (BSP) assay design, methylation-independent BSP, cloning strategy, sequencing and data analysis. Methods that are described nicely elsewhere will not be covered in detail. Instead, the focus will be on tips/tricks and new methods/strategies used in this protocol, including quality control assessment of the DNA before and after bisulfite conversion and a pooled cloning strategy to reduce time, costs and effort during this step. In addition we comment on dealing with bias and improving overall protocol efficiency.

## INTRODUCTION

There are a lot of ways to study DNA methylation. Depending on the scientific question, the samples (type, quality, quantity, number), the laboratory equipment and funds, researchers can compose their most appropriate strategy. Since all methods have their pros and cons, it is vital to evaluate all steps for potential bias, take measures to prevent them and include necessary controls to monitor them [1–2].

Here, we report our strategy for cloning-based locus-specific bisulfite sequencing PCR (BSP) to investigate the methylation status of specific CpG islands at single base resolution. The strategy is partially based on described strategies [3–11], but also contains some useful adaptations. This strategy can be used to investigate if a gene specific expression change in an organism is caused by an altered methylation status of that gene.

First, bisulfite treatment, the gold standard method in DNA methylation studies, will selectively convert “unmethylated” cytosine (C) to uracil (U), while “methylated” C will not be converted [12]. It should be noted that other C-modifications, such as 5-formylcytosine (5-fC) and 5-carboxylcytosine (5-caC), will be converted to U as well, while others, such as 5-hydroxymethylcytosine (5-hmC), will not be converted either. However, adapted methods exist to study these rarer modifications separately [13–14].

Then, PCR is performed to selectively amplify the bisulfite-converted region of interest, whereby U (native C, 5-fC and 5-caC) will be replaced by thymine (T) and non-converted C (native 5-mC and 5-hmC) by C. After Sanger sequencing, all remaining Cs can be considered as “methylated” Cs in the native sequence (5-mC or 5-hmC). We prefer a cloning-based strategy (instead of direct sequencing) in order to obtain DNA methylation haplotypes. In addition, the interpretation of the peaks is unequivocally (no mixed bases, misaligned signals or PCR slippage). In order to make it less laborious, we maximize amplicon lengths based on the bisulfite-converted DNA quality control and use a pooled cloning strategy.

## PROTOCOL

### 1) Sample handling

Because bisulfite sequencing is most successful with intact starting material, tissue/DNA samples should be handled/stored in a way that prevents DNA degradation. Well known key factors are temperature (cold, avoiding freeze-thaw cycles), humidity (dry), sunlight (darkness) and time (quick). For extensive guidelines see [15].

### 2) DNA isolation

Total DNA is isolated with the Quick-DNA Miniprep Plus Kit (including a Proteinase K digest, according to the Zymo Research’s recommendations), described to extract ultra-pure concentrated RNA-free high-quality DNA from a wide range of biological sample types ready for bisulfite sequencing (maximal binding capacity of the column is 25 μg DNA and minimal elution volume is 35 μl). Many other kits or protocols are described that should work equally good [16]. At first use we recommend to isolate DNA from a test sample and evaluate the procedure(s) based on the DNA quality control results (see Protocol, section 3).

### 3) DNA quality control before bisulfite conversion

The quantity and purity of the extracted DNA is measured with Nanodrop as dsDNA (Isogen). Integrity is evaluated by analysing 1 μg of DNA on a 1% agarose gel and by performing the UBC integrity assay on 5 ng DNA (Table 1 and Figure 1.A). The UBC integrity assay consists of a single monoplex PCR reaction amplifying fragments of different lengths (137, 365, 593, 821,… bp) analysed on a 2% agarose gel [17]. Pure and intact DNA will allow amplification of all fragments, while higher degrees of impurity and/or degradation will result in a decrease of amplification products starting with the longer amplicons. Implementation of the multi-use UBC integrity assay in the lab is of particular interest since this single assay can not only be used for quality assessment of DNA from different mammals (checking presence, integrity, amplificability), but also to estimate the DNA contamination level in RNA samples and to perform quality assessment of cDNA reverse transcribed from RNA isolated from any tissue (reflecting the RNA quality). Ideally, DNA should be pure and integer (= OD260/280 around 1.8 on Nanodrop, a high molecular weight band on gel and generation of all amplicons with the UBC integrity assay). Also for this step other methods (e.g. fluorometric- or microfluidic-based methods) can be used to perform DNA quality control [18].

**Table 1.**
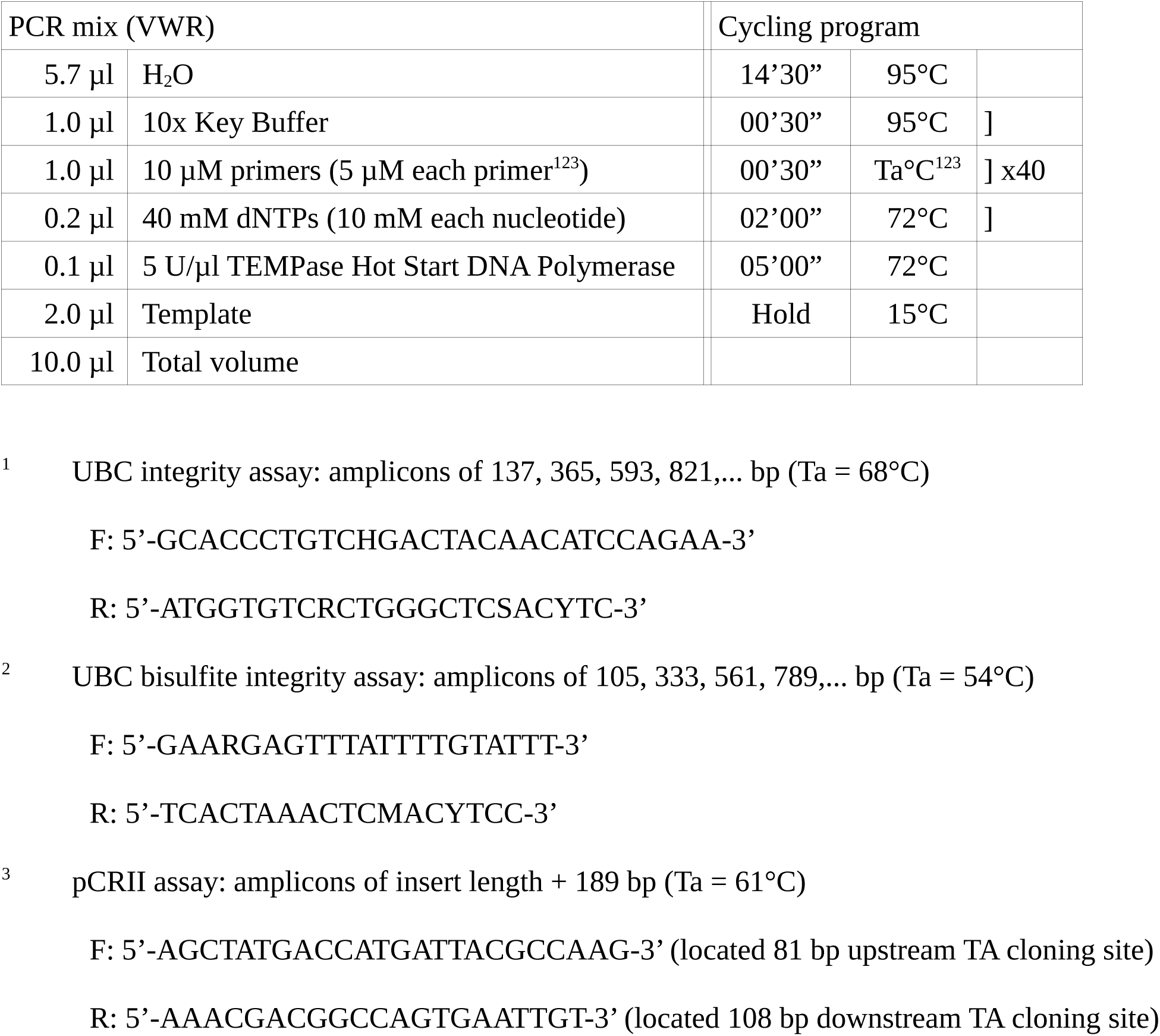
PCR details of the UBC (bisulfite) integrity [17] and pCRII assays.

**Figure 1.**
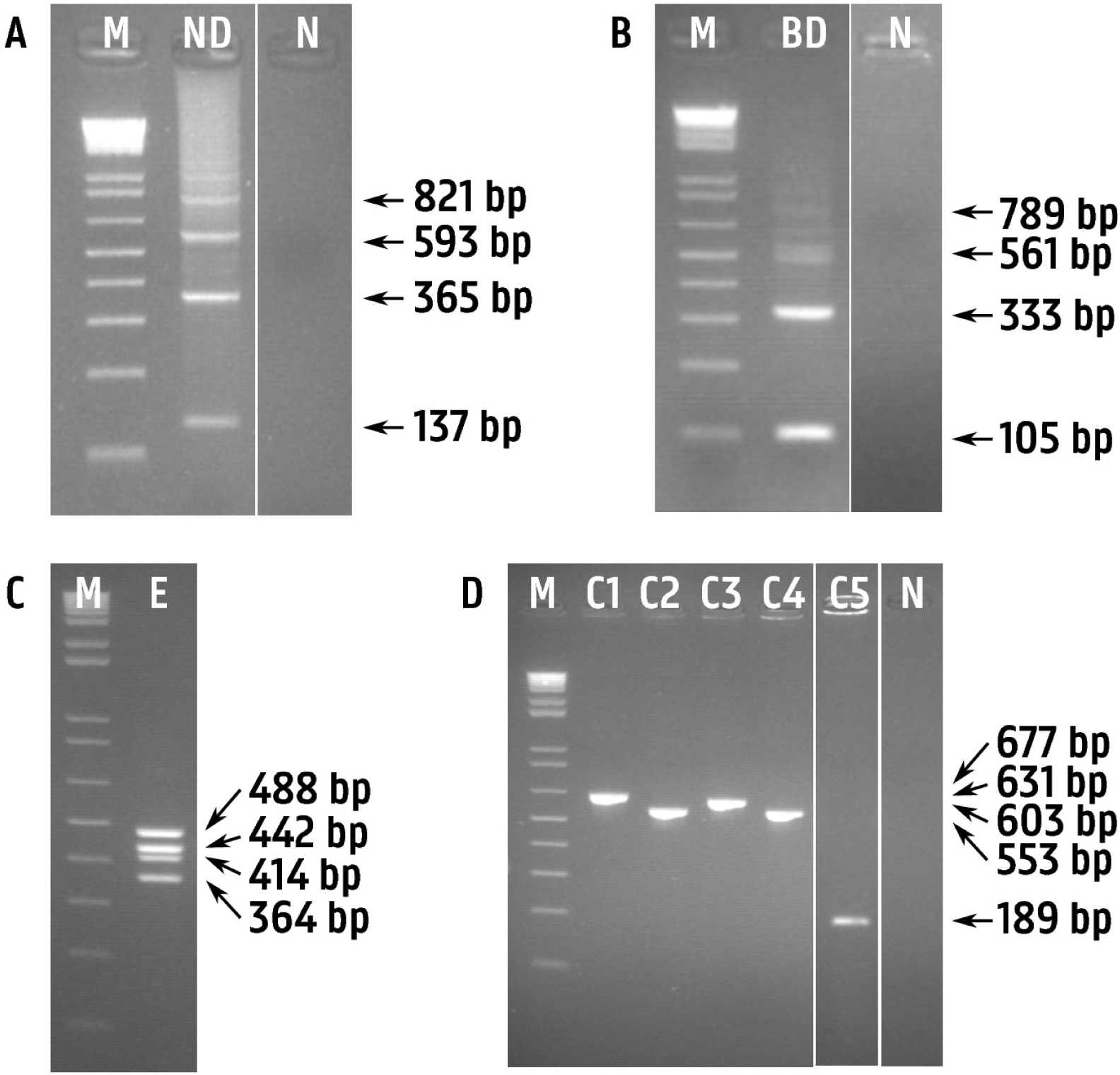
Agarose gels showing A) UBC integrity assay on genomic DNA before bisulfite conversion (Protocol, section 3; adapted from [17]), B) UBC bisulfite integrity assay on genomic DNA from (A) after bisulfite conversion (Protocol, section 5; adapted from [17]), C) eluted 4-amplicon mix amplified on genomic DNA from (B) before cloning (Protocol, section 9; adapted from [33]), and D) colony PCR with pCRII primers on 5 clones containing amplicons from (C) (Protocol, section 9; adapted from [33]). M: 1 kb+ ladder (ThermoFisher Scientific), ND: native DNA, BD: bisulfite-converted DNA, E: eluted 4-amplicon mix, C1-4: clone with an insert, C5: clone without an insert, N: no template control.

### 4) Bisulfite conversion

Bisulfite conversion is performed on 500 ng of RNA-free high-quality DNA with the EZ DNA Methylation-Lightning Kit (according to the Zymo Research’s recommendations), described to convert > 99.5% of unmethylated Cs and to protect > 99.5% of methylated Cs, with a DNA recovery of > 80%. Higher input levels of DNA are not recommended because they might result in incomplete bisulfite conversion. Recommended input levels can go as low as 100 pg, however this will lower proportionally the number of downstream PCR reactions and the maximal fragment length that can be amplified (because of the lower input of damaged DNA, the number of the longer fragments might drop below the threshold for amplification). The bisulfite-converted DNA is eluted in 10 μl (around 40 ng/μl bisulfite-converted DNA). Many other kits or protocols are described that can be used [10,16]. At first use we recommend to perform bisulfite conversion on a test sample and evaluate the procedure(s) based on the DNA quality control results after bisulfite conversion (see Protocol, section 5).

### 5) DNA quality control after bisulfite conversion

The quantity and purity of the DNA after bisulfite conversion, known to damage DNA, is measured with Nanodrop as ssRNA (Isogen). Integrity and amplificability is evaluated by performing the UBC bisulfite integrity assay on 5 ng of bisulfite-converted DNA (Table 1 and Figure 1.B). It is a similar assay as the one used for native DNA, but for DNA after bisulfite conversion [17]. Comparing the results of both integrity assays will give an idea about the impact of bisulfite conversion on the DNA integrity of the sample. It will also give an idea about the maximal fragment length that can be PCR amplified from the sample. Because of the fragility of bisulfite-converted DNA, it is advised to proceed immediately to PCR and freeze the rest in aliquots.

### 6) *In silico* identification of CpG islands

*In silico* identification of CpG islands in target genes is based on common hits in different genome browsers (Ensembl, UCSC and NCBI) and online tools such as CpG Islands (The Sequence Manipulation Suite), DBCAT, Cpgplot (EMBOSS) and MethPrimer [19–25].

### 7) Methylation-independent BSP assay design

Methylation-independent BSP primer design and electronic PCR, detecting potential mispriming sites and undesired PCR products, is performed by BiSearch [26]. To our knowledge, it is the only free software combining BSP primer design and electronic PCR. Customized parameters are discussed below.

Because bisulfite-converted DNA is not complementary anymore, a choice has to be made whether to design primers amplifying the sense or the antisense strand. We suggest to try both strands and choose the most optimal primers. Because of the symmetry of the CpG motifs and the mode of action of the methyltransferases, the methylation status of every CpG motif should be identical to its complement, unless the region of interest is prone to hemimethylation [27]. Signs for hemimethylation can be observed by analysing the methylation status of CpGs in overlapping parts of amplicons targeting the different strands and warrant further investigation.

Amplicon length is based on the length of the CpG island to be analysed (see Protocol, section 6), the integrity and amplificability of the bisulfite-converted DNA (see Protocol, section 5) and the cloning strategy (see Protocol, section 9). Using the described protocol, amplicons up to 800 bp can be amplified starting from high-quality DNA.

Because of the bisulfite conversion, 4-base DNA (25% of A, G, C and T) will be shifted towards 3-base DNA (towards 25% A, 25% G, 0% C and 50 % T), reducing DNA complexity. In order to have the same specificity, bisulfite primers might need to be longer compared to native primers.

In case an estimate of the primer occurrence in a particular template is wanted, the following formula can be used: N × (pA^Na) × (pG^Ng) × (pC^Nc) × (pT^Nt), with N being the number of nucleotides in the template, pA/pG/pC/pT the estimated frequencies of the respective nucleotides in that template (sum should be 1) and Na/Ng/Nc/Nt the number of the respective nucleotides in the primer. In an average mammalian genome of 3×10^9 bp (assuming that every nucleotide appears at 25%), a native primer of 20 bp (containing 5 times each nucleotide) would theoretically occur 0,0027 times (= (3×10^9) × (0.25^5) × (0.25^5) × (0.25^5) × (0.25^5)), so considered to be highly specific. For bisulfite primers in a hypothetical 100% methylated genome (0% C converted to T), it would be the same. In a hypothetical 100% unmethylated genome (100% C converted to T) a similar 20-bp primer (all 5 Cs converted to Ts) would occur 3 times (= (3×10^9) × (0.25^5) × (0.25^5) × (0^0) × (0.5^10)), so considered to be not specific. In a genome where 40% of the Cs would be methylated, a similar 20-bp primer (3 out of 5 Cs would be converted to T) would occur 0.02 times (= (3×10^9) × (0.25^5) × (0.25^5) × (0.1^2) × (0.4^8)), about 10 times less specific than the respective native primer.

Taking into account the completeness of genome databases, the specificity of potential PCR primers can be checked via the fast PCR tool of BiSearch using the 16-mer mismatch string parameter to specify nucleotide specific differences (e.g. random mismatch in the genome and Cs that might or might not be converted after bisulfite treatment). In addition, the native versions of the bisulfite primers can be checked for known SNPs via NCBI-BLAST in order to prevent null-alleles [28].

To avoid that primer annealing is affected by the methylation status of the primer target sequence, primers should not contain CpGs. In case they do, degenerate primers should be designed with a Y (C or T) instead of a C. Amplifying unconverted DNA can be prevented by including some non-CpG Cs in the native primer sequence (they will be replaced by Ts in the bisulfite primer and as a result only be specific for converted DNA). To make sure, it can be experimentally verified that methylation-independent primers do not amplify unconverted DNA.

Annealing temperatures should be as high as possible to prevent potential secondary structures in the template and avoiding primer dimer formation. Inter-primer melting temperature (Tm) differences should be as low as possible (lower than 1°C) to prevent non-binding of the primer with the lower Tm or non-specific binding of the primer with the higher Tm.

### 8) Methylation-independent BSP

Because PCR on bisulfite-converted DNA is prone to non-specific amplification due to its high AT content, it is strongly recommended to use a HotStart polymerase. From the wide range of available DNA polymerases, we use TEMPase HotStart Polymerase (according to VWR’s recommendations), designed to diminish the formation of non-specific priming events during reaction set-up and the first ramp of thermal cycling. It is a non-proofreading DNA polymerase (produces 3’-A overhangs), allowing TA cloning (see Protocol, section 9). Other DNA polymerases can be used, but not all. Archaeal polymerases, such as the high-fidelity polymerases Vent and Pfu, are unable to efficiently copy bisulfite-converted DNA due to the stalling triggered by template uracil [29]. In addition, unmodified high-fidelity polymerases will complicate subsequent TA-cloning, because they do not produce 3’-A overhangs.

PCR is performed for 40 cycles (30”-95°C, 30”-Ta, 2’-72°C) with 5 ng bisulfite-converted DNA as a template (= on average 80 reactions can be performed per conversion) on a S1000 Thermal Cycler (Bio-Rad) with gradient function. During optimization of the assays, a 5-point gradient PCR is performed with as annealing temperature (Ta) the predicted Tm −4°C, −2°C, +0°C, +2°C and +4°C. Amplicons are analysed on a 2% agarose gel. The averaged Ta of all Ta with specific amplification is chosen as assay Ta. Because of the complexity of the PCR reaction (fragmented DNA, low complexity target, presence of U) it might be needed to increase extension times.

### 9) Cloning strategy

If multiple fragments need to be analysed, we opt for a pooled cloning strategy in pCRII (TA-cloning kit, Invitrogen) in order to reduce time, costs and effort during this step. Ideally, pooled amplicons should differ in length (the longer the amplicons, the longer the difference). After PCR on bisulfite-converted DNA with a non-proofreading DNA polymerase, the different amplicons are analysed on a 2% agarose gel, cut out with a scalpel and eluted together (up to 4 different amplicons) with the GENECLEAN II kit (MP Biomedicals) in 8 μl. One μl of the eluted amplicon mix is analysed on a 2% agarose gel to validate the amplicon quantities (Figure 1.C). Six μl of the eluted amplicon mix is then ligated in 1 μl pCRII with 1 μl T4 DNA ligase (= 1 U) and 2 μl 5x T4 DNA ligase buffer at 14°C overnight. Two μl of the ligation mix is then transformed into 50 μl Subcloning Efficiency DH5α Competent Cells and grown overnight on LB plates containing 100 μg/ml ampicillin and 50 μg/ml X-gal (allowing blue/white screening; according to Invitrogen’s instructions). The next day, individual white colonies (containing 1 insert) are striped on new plates and grown overnight. The next day, a tip-point of cells is resuspended in 100 μl water and 2 μl is used for colony PCR. If the amplicon length difference of the pooled fragments can be distinguished on a 2% agarose gel, pCRII primers bordering the TA cloning site can be used to amplify the insert to be sequenced (Table 1). Two μl of the PCR product can be analysed on a 2% agarose gel to check the amount and the identity of the insert based on fragment length (Figure 1.D). By doing so, the amount of input for sequencing and the number of clones to be sequenced from each pooled fragment can be controlled. If some of the pooled fragments can not be distinguished from each other by length, they can be first cut with a specific restriction enzyme before gel analysis, or fragment specific primers can be used on the undetermined clones. If preferring another cloning strategy, the above-mentioned issues can be adopted as needed.

### 10) Sequencing

The rest of the colony PCR product (= 8 μl) of the selected clones (at least 6 for every fragment) is cleaned-up for Sanger sequencing by adding 4 U exonuclease I (Bioké) and 2 U antarctic phosphatase (Bioké), and incubating for 30 min at 37°C (enzymatic reaction) and 15 min at 80°C (enzyme inactivation). Two μl of the treated PCR product is usually (depending on its amount based on Figure 1.D) used for the sequencing reaction with the BigDye Terminator v3.1 Cycle Sequencing Kit (Applied Biosystems; Table 2) using one or both (depending on the length of the insert) PCR primers as individual sequencing primer.

**Table 2.** Sanger sequencing details (BigDye Terminator v3.1 Cycle Sequencing Kit, Applied Biosystems)

| Sequencing mix |  | Cycling program |  |  |
| --- | --- | --- | --- | --- |
| 3.0 µl | H<sub>2</sub>O | 2'00" | 95°C |  |
| 0.5 µl | Ready Reaction mix | 0'20" | 95°C | ] |
| 2.0 µl | 5x sequencing buffer<sup>1</sup> | 0'10" | 60°C | ] x30 |
| 1.0 µl | GC-rich solution (Roche) | 4'00" | 65°C | ] |
| 1.5 µl | Sequencing primer (2 µM) |  |  |  |
| 2.0 µl | Template |  |  |  |
| 10.0 µl | Total volume |  |  |  |

<sup>1</sup> 200 mM Tris-HCl, pH 8 + 5 mM MgCl<sub>2</sub>

### 11) Data analysis

The chromatograms are inspected manually for errors and the sequences are trimmed (insert without amplicon specific primer sequences, because they do not represent the methylation status of the native fragment) with BioEdit (free software) [30]. Extracting the methylation data (including quality control and visualisation in lollipop-style) is performed with BiQ Analyzer (free software) [31].

## COMMENTARY

### 1) Dealing with bias

It is important to evaluate every step of the protocol for a potential introduction of bias. Most critical is probably the conversion efficiency of the bisulfite treatment. According to the specifications of the kit used in our protocol the conversion efficiency is > 99.5% (= less than 1 error per 200 CpGs). For a hypothetical amplicon of 400 bp containing 40 CpGs, this would mean less than 1 CpG error per 5 amplicons. Experimental bisulfite conversion efficiencies can be estimated by calculating the percentage of non-CpG Cs in the native amplicon sequence that are really converted to Ts in the bisulfite-converted sequence (one of the QC parameters of BiQ Analyzer). Including non-CpG Cs in the native primer sequence will prevent amplification of unconverted DNA and thus lower potential bias.

Another source of potential bias is caused by PCR. According to the PCR Fidelity Calculator (ThermoFischer Scientific) [32], amplification of the hypothetical 400-bp fragment for 40 cycles with *Taq* DNA polymerase would introduce 1 error in 1/3 of the amplicons. Because only C>T errors at methylated CpGs or T>C errors at unmethylated CpGs (= 1/3 of all possible errors) of the 40 CpGs of the amplicon (= 1/10 of the sequence) would create bias (all other errors would be noticed as errors), this would theoretically result in a wrong determination of the methylation status of only 1 CpG per 90 amplicons (= 1/3 * 1/3 * 1/10; almost 20-fold less than bisulfite conversion errors). Because we perform cloning-based sequencing involving colony PCR with a non-high-fidelity DNA polymerase, a similar PCR bias is created during this step. However, there would be no implications here when using a high-fidelity DNA polymerase to lower this bias. In addition, it might even lower potential PCR slippage (another PCR bias), typically due to sequential Ts (N>9). In case PCR slippage during colony PCR hinders sequencing (not an issue before cloning), sequencing could be performed on DNA extracted from a single clone (instead of performing colony PCR).

To test if the primers amplify methylation independent (and not in favour of unmethylated templates), PCR on a 50:50 methylated/unmethylated bisulfite converted control sample is frequently performed. To make sure that all tested amplicons are really 50:50 methylated/unmethylated, all regions under investigation are first PCR amplified with native primers on native DNA as a template (these amplicons can contain multiple overlapping BSP amplicons). These 100% unmethylated amplicons are mixed and split into two parts. One part will serve as the unmethylated part, the second part will be 100% CpG methylated by a CpG methyltransferase treatment (M.SssI, Bioké). This can be verified by cutting an aliquot of both parts with *Hpa*II (Bioké). It will cut unmethylated CCGG (= unmethylated part 1), but not methylated CCGG (= methylated part 2). Both parts are then mixed, cleaned-up (QIAquick DNA purification kit, Qiagen) and bisulfite converted. Then all BSP assays are performed and amplicons digested with *Hpa*II and *Taq*I. *Hpa*II (cuts CCGG) will not cut bisulfite converted DNA (unmethylated CCGG will be converted to TTGG and methylated CCGG will be converted to TCGG). So, if none of the amplicons are digested it means that the bisulfite conversion was successful. *Taq*I (cuts TCGA) will not cut the unmethylated part (all native TCGA sequences are converted to TTGA), but will cut the methylated part (all native TCGA will not be converted and all CCGA will be converted to TCGA). So, if half of the amplicons are digested it means that the assays amplify methylation independent.

In order to have a reliable estimate of the methylation status and to minimize the effect of a potential error at every single CpG, six clones are sequenced. To obtain a more precise determination of the methylation status of partially methylated loci, additional clones containing those loci can be sequenced or methylation specific primers targeting those loci can be used as deemed fit. In case of doubt about cloning bias, direct sequencing can be performed and the results compared with the ones obtained via cloning-based sequencing.

Results from identical sequences from overlapping amplicons can also be used to evaluate the reliability of the results. In addition, chromatograms are inspected manually in order to avoid base calling errors during sequencing.

Finally, it is obvious that positive and negative controls should be performed and contamination should be avoided at any time.

### 2) Improving overall protocol efficiency

It is important to avoid DNA degradation because bisulfite sequencing is most successful with intact starting material. In addition, it will allow you to amplify longer amplicons (determined by the UBC bisulfite integrity assay), resulting in less amplicons to process. In order to maximize the chance to reach the threshold number of fragments for amplification of these longer fragments, the maximal advised DNA input for bisulfite conversion is used.

The protocol involves 2 PCR steps, one PCR on bisulfite-converted DNA before cloning and one colony PCR. Because the theoretical bias created by the DNA polymerase is about 20-fold lower than the bias created during bisulfite conversion, there is no big benefit to use more expensive high-fidelity DNA polymerases, that even might complicate TA cloning. In our opinion, if primer design guidelines are followed properly, there is no need for optimization (except for determining the optimal experimental Ta) or performing (semi)-nested PCR.

Because cloning-based bisulfite sequencing is labour intensive, the pooled cloning strategy really makes it more efficient. If pooled amplicons differ at least 50 bp in length, their clones can easily be distinguished from each other by a single colony PCR with universal vector primers. If not, additional work might be needed to identify the clones in order not to sequence too many clones containing the same amplicon. Although we were able to amplify amplicons of 800 bp, it is always easier to amplify, clone and sequence smaller amplicons. In the pooled cloning strategy we used no more than 4 amplicons between 350 and 500 bp.

At last, using free data analysis software, such as BiQ Analyser, minimizes errors and speeds up the analysis.

## ACKNOWLEDGEMENTS

We wish to thank Carolien Rogiers, Dominique Vander Donckt, Linda Impe and Ruben Van Gansbeke for excellent technical assistance.

